# Global Cluster Analysis and Network Visualization in Chordoma: A Scientometric Mapping

**DOI:** 10.1101/2023.11.19.567758

**Authors:** Bo-Wen Zheng, Bo-Yv Zheng, Hua-Qing Niu, Ming-Xiang Zou, Fu-Sheng Liu

## Abstract

**Background:** A recent scientific and comprehensive examination of the present status and future directions in chordoma research is a subject worth exploring. This study endeavors to investigate and depict the quantitative and qualitative development, research frontiers, and future trends in chordoma using bibliometric approaches.

**Methods:** A total of 2510 publications from the Science Citation Index Expanded of Web of Science Core Collection were obtained from 1991 to 2021 and selected for bibliometric analysis. Bibliometric and knowledge-map analysis were performed using visualized approaches such as VOS viewer and CiteSpace, along with Origin 8.0 and GraphPad prism 8 tools.

**Results:** The top five countries with the highest number of publications are the United States, China, Japan, Italy, and Germany. The most impactful journal is *World Neurosurgery*, while the most cited journal is *Journal of Neurosurgery*. The five most common research fields are neuroscience neurology, surgery, oncology, pathology, and radiology nuclear medicine medical imaging. Harvard University has the highest number of publications among institutions, and Gokaslan, Ziya L. has published the most papers and has the highest total link strength. The article “Chordoma: incidence and survival patterns in the United States, 1973-1995” has the most citations. The co-occurrence analysis includes 744 keywords that were categorized into five clusters: radiotherapy study, basic and clinical trill research, diagnostic study, surgical approach research, and prognostic study. CiteSpace identified 12 clusters of keywords, with clusters 0-5, 7, and 12 being former hotspots, clusters 6 and 11 being mid-period hotspots, and cluster 8 being the current research hotspot.

**Conclusions:** This bibliometric analysis and visualized study of chordoma’s global research field in the past 30 years is the first of its kind. The findings may help academic researchers comprehend chordoma’s dynamic evolution and research trends.

Investigating the current status and future trends in chordoma research is a valuable undertaking. Using a bibliometric approach, this study aims to quantitatively and qualitatively explore the development and frontiers of chordoma research. A total of 2510 publications were analyzed, downloaded from the Science Citation Index Expanded of Web of Science Core Collection between 1991 and 2021. Bibliometric and knowledge-map analysis were performed using visualized approaches, Origin 8.0, and GraphPad prism 8 tools. The top five countries with the most publications were the USA, China, Japan, Italy, and Germany. World Neurosurgery was the most influential journal, and the Journal of Neurosurgery was the most co-cited. The research fields focused on neuroscience, neurology, surgery, oncology, pathology, and radiology nuclear medicine medical imaging. Harvard University produced the highest number of publications, and Gokaslan, Ziya L had the most papers and total link strength. The co-occurrence analysis included 744 keywords and was classified into five clusters: radiotherapy study, basic and clinical trill research, diagnostic study, surgical approach research, and prognostic study. Additionally, 12 clusters of keywords were identified by CiteSpace, with cluster 0 to 5, cluster 7, and cluster 12 being former appeared hotspots, clusters 6 and 11 being mid-period appeared hotspots, and cluster 8 being present research hotspots. This is the first comprehensive bibliometric analysis and visualized study of the global chordoma research field in the past 30 years, providing insights into its dynamic evolution and research trends. It may be beneficial for researchers to gain a better understanding of chordoma research.

**Level of Evidence:** Level 3

**Short Summary:** This investigation represents the inaugural comprehensive and scientific bibliometric analysis and visualization of the global chordoma research landscape over the preceding three decades. The present inquiry offers advantages to scholarly investigators seeking to gain a nuanced understanding of the dynamic development and research tendencies pertaining to chordoma.

## Introduction

Chordoma is an exceedingly rare and malignant neoplasm that arises from vestigial notochord remnants, typically in the sacrum or skull base, affecting patients of all ages[1, 2]. Normally, notochordal cells differentiate into the nucleus pulposus of the intervertebral discs, but a subset of these cells may be distributed within the vertebral bodies through an incompletely understood mechanism, and this subset, called the notochordal remnants, can undergo malignant transformation into chordoma[3]. These infrequent tumors have an annual incidence of approximately 1 per 1 million people/year and exhibit local aggressiveness with a high recurrence rate[4, 5]. Although chordomas can occur at any spinal level, they are primarily found in the sacrococcygeal region (50-60%) and often involve the clivus at the skull base (30%)[6]. Males are more commonly affected, with an average age at diagnosis of approximately 60 years. However, presentation at the skull base is gradually affecting younger populations and children[5]. In sacrococcygeal and vertebral body cases, there is a male predominance (2:1), while tumors affecting the skull base do not show any sex difference[7].

Veronica Ulici described chordoma as a lobulated and expansile intraosseous mass that initially penetrates the bony cortex and subsequently invades adjacent tissues[8]. However, the pathogenesis of chordoma remains unclear to date. The tumor cells of chordoma are usually characterized by notochordal differentiation, confirmed by the overexpression of brachyury (a nuclear transcription factor involved in notochordal growth regulation)[3]. The World Health Organization categorizes chordoma into three types: conventional chordoma, poorly differentiated chordoma, and dedifferentiated chordoma[9]. Chordoma may express specific keratins (such as cytokeratin 8 [CK8], CK18, CK19) and is typically positive for S100 protein and epithelial membrane antigen (EMA) while being negative for CK7 and CK20[10–12]. Differential diagnosis of the various benign and malignant entities of chordoma is possible. For example, benign lesions with morphologic similarity include benign notochordal cell tumors (BNCTs) and ecchymosis physaliphora (EP).

The treatment of chordoma primarily relies on surgical excision, which is often incomplete and yields poor results[13]. However, several molecular alterations that have been identified with reasonable accuracy have led to ongoing clinical trials using targeted therapy. En bloc resection is currently the mainstay treatment for chordoma[13]. However, the axial location of most cases makes complete tumor resection challenging. Radiation therapy is suitable for most unresectable patients, although adjuvant radiation may be appropriate in some cases. Unresected chordomas underwent high-dose photon/proton radiation and displayed a 5-year local control rate of approximately 85%, with an approximately 20% rate of distant failure and approximately 89% disease-specific survival[8]. Conventional chemotherapy is generally ineffective against chordoma; hence, clinical trials are currently investigating the use of targeted therapy[8]. Tazemetostat (a small-molecule inhibitor of enhancing zeste homolog 2, EZH2) and brachyury therapeutic vaccines have been evaluated in numerous clinical trials[8, 14].

Based on metrology characteristics and databases in the literature, we conducted a bibliometric analysis to objectively and visibly elucidate the development trends of research activities[15]. Bibliometric analysis, a quantitative measure that analyzes bibliographic data in information and library science, can forecast the evolution of a particular field[16]. Furthermore, it enables identification of the contributions of authors, institutions, countries, and journals in the field, as well as evaluation of collaborative relationships among them[17]. Notably, the bibliometric evaluation is significantly influenced by the period of the selected studies, emphasizing the importance of timely updating of scientific research to obtain the research frontiers[18]. Additionally, the analysis of frequently occurring keywords in publications provides supportive evidence for future trends[19]. These viable methods have been applied to evaluate research trends in various fields, including orthopedics, cancers, coronavirus, and others[20–24]. However, the global development trends of chordoma research remain underexplored. To address this gap, we conducted a bibliometric analysis of chordoma research studies from 1991 to 2021 to examine current status and global trends. The analysis encompassed the number of countries, publications, institutions, journals, authors, and keywords to summarize developmental trends and hot topics, providing guidance for future research.

## Materials and methods

### Data source and search strategy

The present study relied on the Web of Science (WoS), an authoritative database platform with over 12,000 academic journals, to acquire global academic information[25]. Bibliometric analysis was employed to analyze data relevant to chordoma research, following previous studies[18]. To avoid bias, all publications were collected from SCI-Expanded of WoSCC, and the database expiration date was fixed at 15 June 2022 due to the rapid database renewal. The search strategy was designed as follows: theme = Chordoma AND Language = (English) AND publishing year = (1991-2021) AND Document types = (ARTICLE OR REVIEW). Furthermore, specific countries/regions were indexed in WoSCC for more refined information.

### Data Collection and Refinement

The recorded information of published papers, including titles, authors’ names, nationalities, affiliations, the year of publication, name of publishing journals, abstract, and keywords, were downloaded and saved as .txt files from the SCIE database of WoSCC and imported into Excel 2021. Two coauthors manually screened the publications related to chordoma research, and all irrelevant publications were filtered and resolved through discussions with two experts to decide their inclusion in the present study. Finally, all coauthors independently cleaned and analyzed the data using Excel 2021, Origin 8.0, and GraphPad Prism 8.

Figure 1 depicts the publishing criteria, namely: (1) documents focused on the theme of chordoma research; (2) document types were restricted to articles or reviews; and (3) papers were written in English. The exclusion criteria were defined as follows: (1) themes unrelated to chordoma research; and (2) publications such as comments, news, briefings, and meeting abstracts, among others.

**Figure 1.**
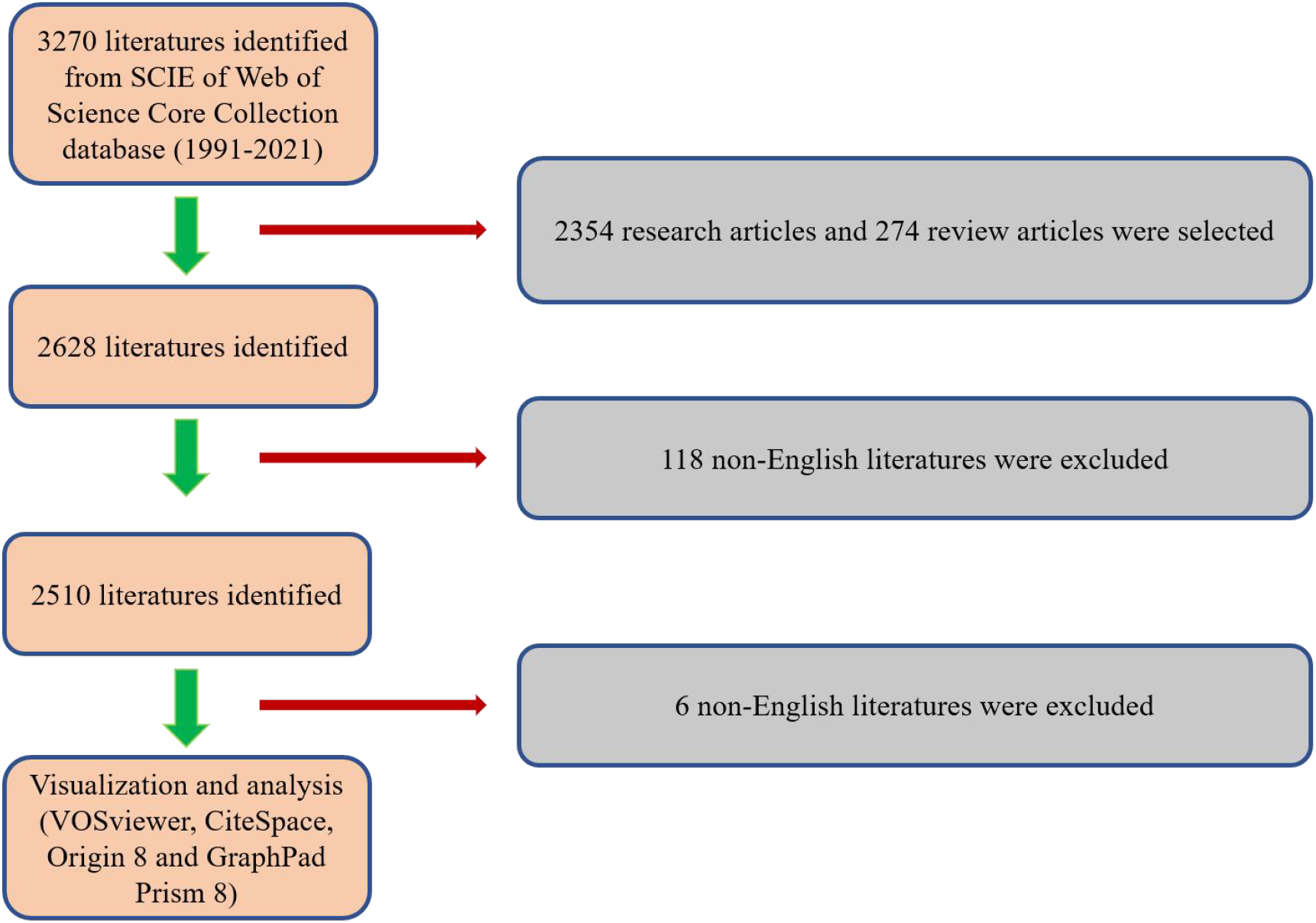
Flowchart of the screening process.

### Bibliometric Analysis

Given its intrinsic function in identifying the fundamental features of collected publications, the SCIE database from WoSCC was the optimal choice for conducting bibliometric analysis. The total number of publications and relative research interest (RRI), defined as the number of publications in one field by all field publications per year, were analyzed using GraphPad Prism 8. To generate the world map of publications, we used R software, which includes python + numpy + scipy + matplotlib[16]. The numbers of publications of different countries/regions were analyzed using Origin 8 software. Subsequently, the time curve of publications was plotted based on a previous study[16]. Total citations and average citations per item of different countries/regions were analyzed using Origin 8.0. The H-index, which indicates that a researcher or country has published H papers and that they have been cited at least H times, was used as a crucial indicator of article quality evaluation. It was selected to assess research’s academic productivity and influence, and it also reflects the number of publications and corresponding citations. Furthermore, the top 25 journals, research orientations, authors, institutions, and funds were analyzed using Origin 8 software.

### Visualized Analysis

We employed VOSviewer (Leiden University, Leiden, the Netherlands) software to visualize and illustrate bibliometric networks, providing more comprehensive information on the cocitation analysis, coauthorship analysis, and co-occurrence analysis in our present study[26]. Specifically, VOSviewer was primarily utilized for bibliographic coupling, cocitation, coauthorship, and co-occurrence analyses.

## Results

### Global contribution to the research of chordoma

In the current investigation, 3270 publications were initially identified from the WoSCC’s SCIE database covering the period 1991-2021. Following the screening process, 2504 articles that conformed to the search criteria were selected for further analysis (Figure 1). The data depicted in Figure 2A indicates a positive trend in the annual number of publications related to chordoma research from 1991 to 2021. The number of publications reached a peak of 190 in 2021, constituting 7.57% of the total corpus. A total of 63 countries and regions made noteworthy contributions to this research domain. Figure 2B and **2C** reveal that the United States made the most significant contribution with 1000 articles (39.84%), followed by China (333, 13.27%), Japan (256, 10.20%), Italy (211, 8.41%), and Germany (178, 7.09%). It is noteworthy that China, Japan, Italy, and Germany collectively contributed to 33.3%, 25.6%, 21.1%, 17.8%, and 15.4% of the US publications, respectively (Figure 2C). In general, our statistical analysis indicates that China has witnessed a rapid surge in publication growth since 2013 (Figure 2D), which may be attributed to its economic expansion. We predict a continuous upward trend in chordoma-related publications from China in the near future. The logistic regression model applied to the publication growth curve predicts that the number of publications in chordoma research will increase from 34 in 1991 to about 381 by 2031 and 1000 by 2044 (Figure 2E). The United States (34354), Italy (6672), Japan (5781), Germany (5522), and England (4521) emerged as the top 5 countries in terms of total citation frequency (Figure 3A). Before Switzerland (40.68), England (39.36), Canada (34.86), the USA (34.35), and Sweden (42.36) had the highest average citation frequencies (Figure 3B). In terms of the H-index, the USA ranked first with 91, followed by Italy (45), Japan (43), Germany (41), and England (40) (Figure 3C).

**Figure 2.**
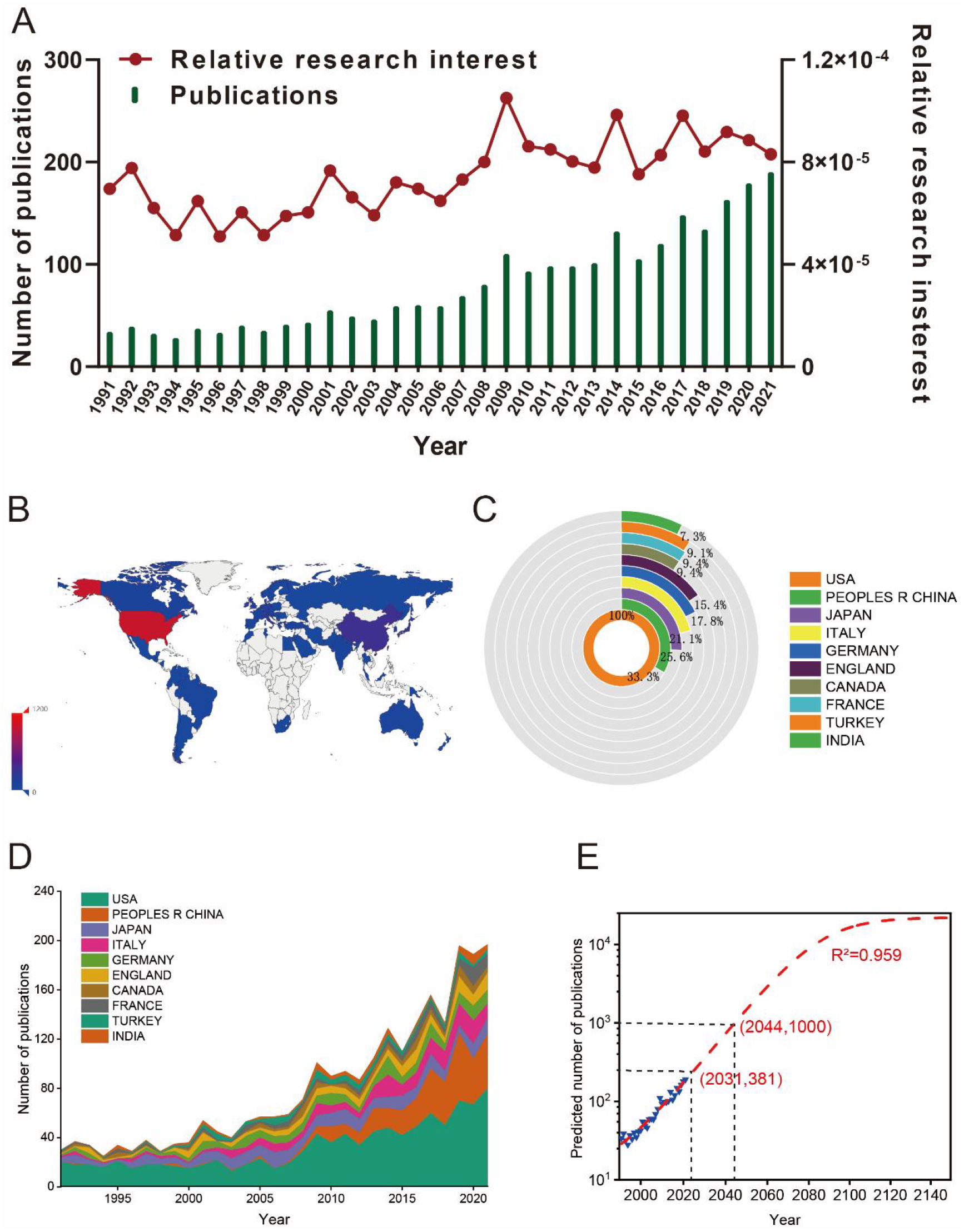
(A) The global number (blue bars) and relative research interests (red curve) of publications related to chordoma research from 1991-2021. (B) Distribution of chordoma research from 1991-2021 in world map. (C) The sum of publications chordoma research from the top 10 countries and regions. (D) The annual number of publications in the top 10 most productive countries from 1991 to 2021. (E) Model fitting curves of global trends in publications related to chordoma research per year (R^2^=0.959, (2031,381) indicates that the total publications will up to 381 in year of 2031), (2044,100) indicates that the total publications will up to 1000 in year of 2044).

**Figure 3.**
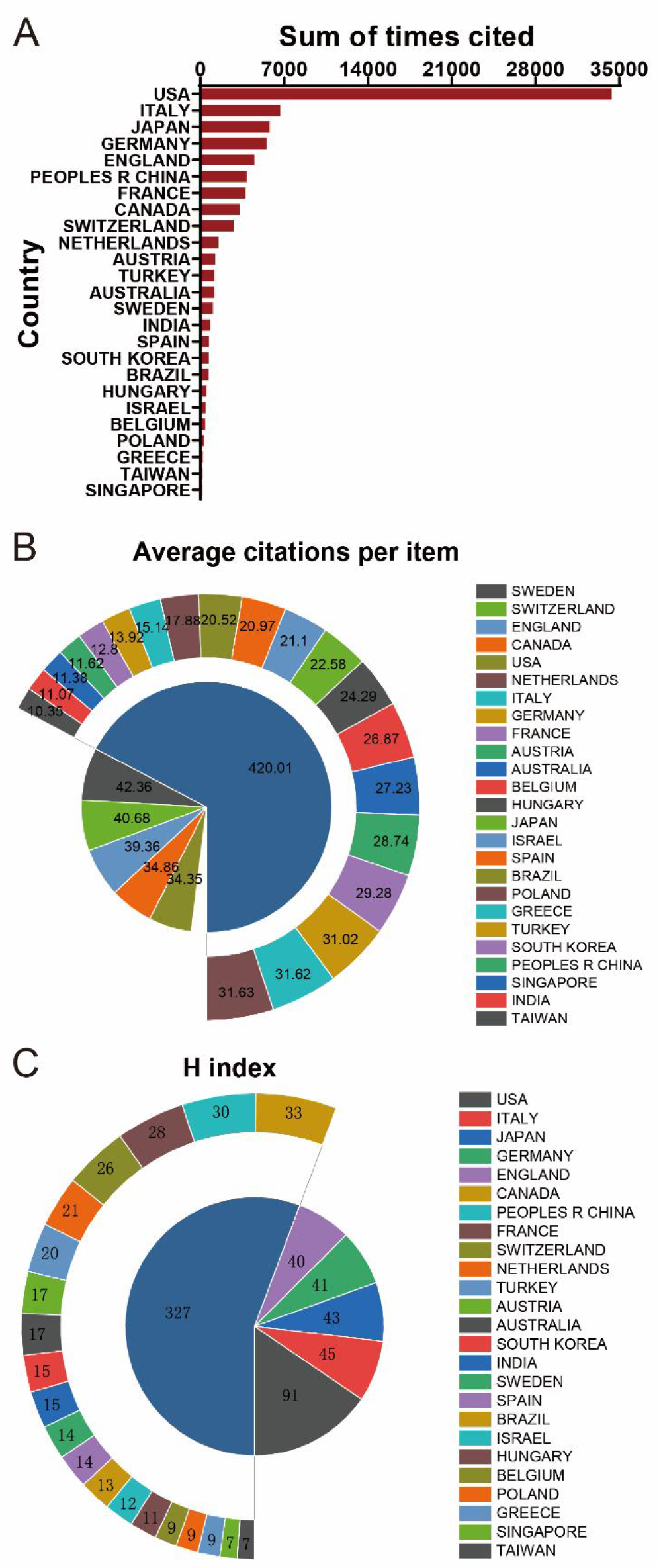
(A) The top 25 countries/regions of total citations related to chordoma research from 1991-2021. (B) The top 25 countries/regions of the average citations per publication related to chordoma research from 1991-2021. (C) The top 25 countries/regions of the publication H-index related to chordoma research from 1991-2021.

### Analysis of Global Publications

Initially, a comprehensive journal distribution analysis was performed to investigate the prominent journals in this field, wherein the top 10 journals with the most publications were tabulated in **Table 1**. The foremost journal, World Neurosurgery (impact factor = 2.210, 2021), exhibited 96 publications, followed closely by Neurosurgery (IF = 5.315, 2021) with 89 publications, Journal of Neurosurgery (IF = 5.408, 2021) with 86 publications, Spine (IF = 3.241, 2021) with 68 publications, and International Journal of Radiation Oncology Biology Physics (IF = 8.013, 2021) with 66 articles.Furthermore, the top 10 research orientations related to chordoma research are displayed in **Table 2** for research orientation analysis, highlighting the prevalent research fields of neuroscience neurology, surgery, oncology, pathology, and radiology nuclear medicine medical imaging.

**Table 1.** The top 10 productive journals related to chordoma research from 1991-2021.

| **Rank** | **Journal** | **Article counts** | **Percentage%** | **IF**<br>**(2021)** |
| --- | --- | --- | --- | --- |
| 1 | *World Neurosurgery* | 96 | 3.825 | 2.816 |
| 2 | *Neurosurgery* | 89 | 3.546 | 5.315 |
| 3 | *Journal of Neurosurgery* | 86 | 3.426 | 5.408 |
| 4 | *Spine* | 68 | 2.709 | 3.241 |
| 5 | *International Journal of Radiation*<br>*Oncology Biology Physics* | 66 | 2.629 | 6.686 |
| 6 | *European Spine Journal* | 53 | 2.112 | 2.721 |
| 7 | *Journal of Neurosurgery Spine* | 53 | 2.112 | 3.467 |
| 8 | *Acta Neurochirurgica* | 49 | 1.952 | 2.816 |
| 7 | *Journal of Clinical Neuroscience* | 36 | 1.434 | 2.116 |
| 10 | *Journal of Neuro Oncology* | 36 | 1.434 | 4.506 |

**Table 2.**
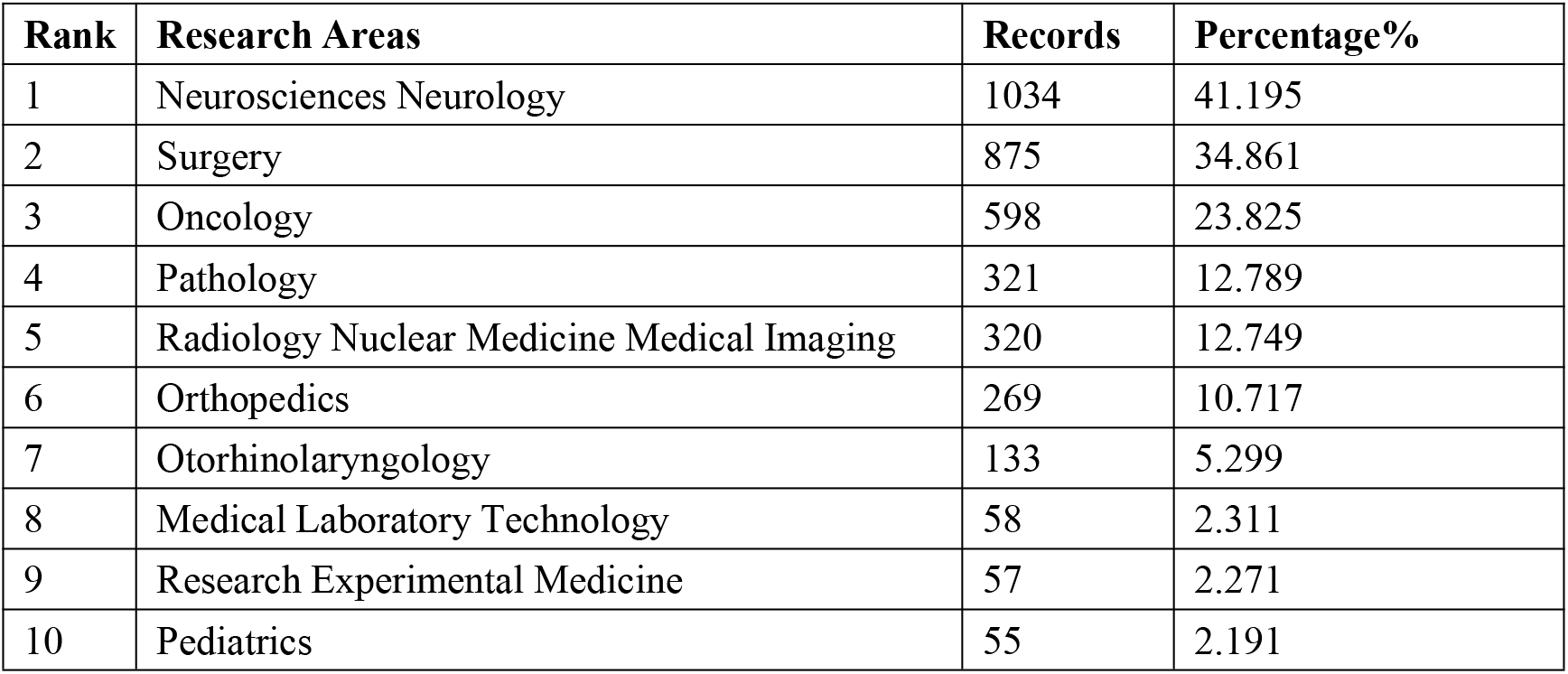
The top 10 well-represented research areas related to chordoma research from 1991-2021.

As for leading author analysis, the top 10 authors contributed a total of 710 publications, encompassing 28.29% of all publications in this field of chordoma research. **Table 3** reveals that Gokaslan, ZL was the leading author with 58 publications, followed by Hornicek, FJ with 45 publications, Debus, J with 39 publications, Sciubba, DM with 38 publications, and Boriani, S with 36 publications. Concerning the publication ranking of institutional output, **Table 4** provides the top 10 contributing institutions, wherein Harvard University claimed the top spot with 151 publications, trailed by the League of European Research Universities Leru with 150 publications, while Massachusetts General Hospital attained the third rank with 121 publications. The top 10 funding sources are presented in **Table 5**, indicating that the National Institutes of Health (NIH, USA) and the United States Department of Health and Human Services provided funding for 148 publications, while the National Natural Science Foundation of China (NSFC) funded 110 publications, and Nih National Cancer Institute NCI funded 106 publications. The Chordoma Foundation was ranked fifth, providing funding for 26 publications.

**Table 3.** The top 10 authors with the most publications related to chordoma research from 1991-2021.

| Rank | High Published Authors | Country | Article counts | Percentage % |
| --- | --- | --- | --- | --- |
| 1 | Gokaslan ZL | Japan | 58 | 2.311 |
| 2 | Hornicek FJ | USA | 45 | 1.793 |
| 3 | Debus J | Germany | 39 | 1.554 |
| 4 | Sciubba DM | USA | 38 | 1.514 |
| 5 | Boriani S | Italy | 36 | 1.434 |
| 6 | Schwab JH | USA | 34 | 1.355 |
| 7 | Gardner PA | USA | 30 | 1.195 |
| 8 | Wu Z | China | 30 | 1.195 |
| 9 | Rhines LD | USA | 28 | 1.116 |
| 10 | Snyderman CH | USA | 28 | 1.116 |

**Table 4.**
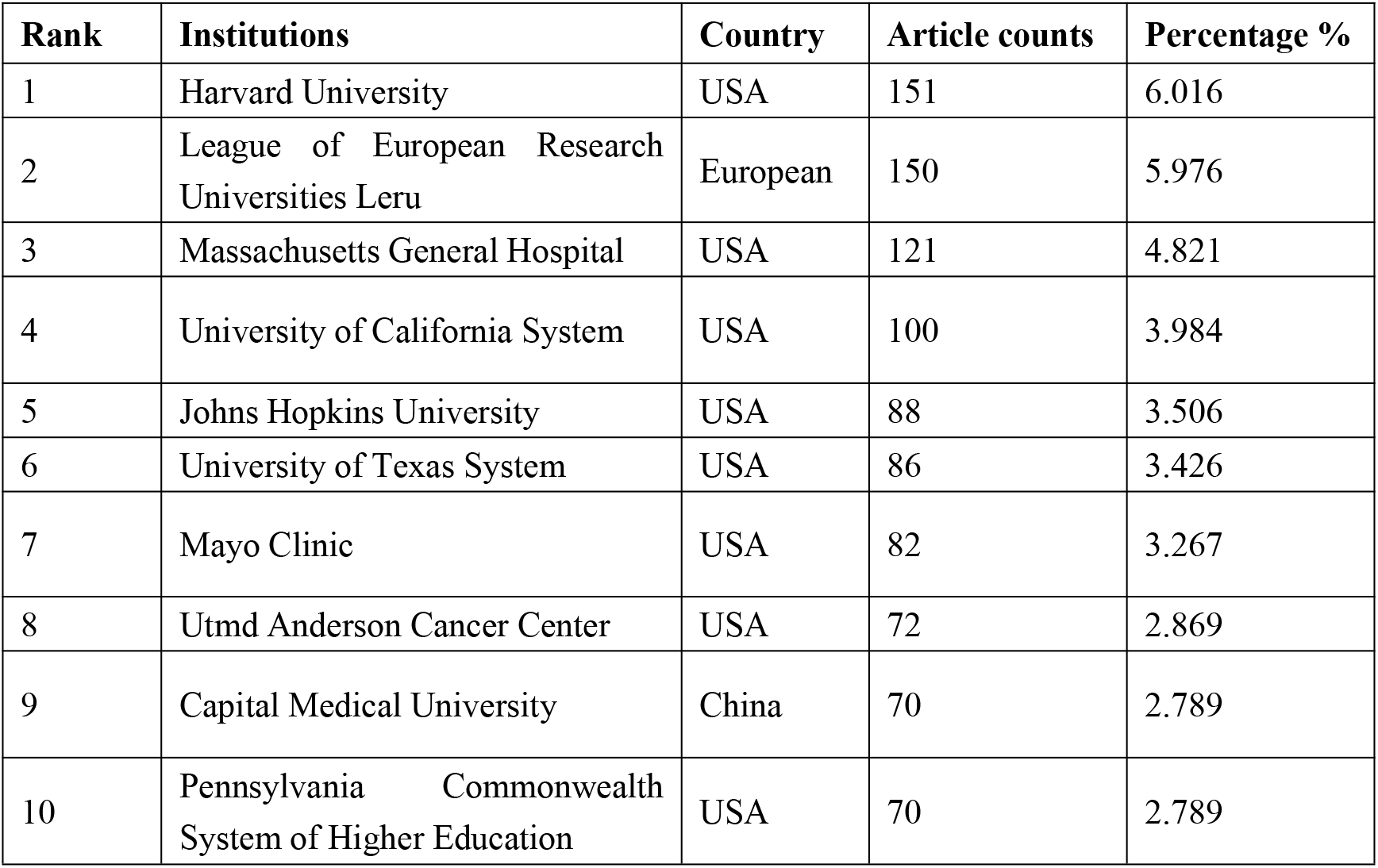
The top 10 institutions with most publications related to chordoma research from 1991-2021.

| Rank | Institutions | Country | Article counts | Percentage % |
| --- | --- | --- | --- | --- |
| 1 | Harvard University | USA | 151 | 6.016 |
| 2 | League of European Research Universities Leru | European | 150 | 5.976 |
| 3 | Massachusetts General Hospital | USA | 121 | 4.821 |
| 4 | University of California System | USA | 100 | 3.984 |
| 5 | Johns Hopkins University | USA | 88 | 3.506 |
| 6 | University of Texas System | USA | 86 | 3.426 |
| 7 | Mayo Clinic | USA | 82 | 3.267 |
| 8 | Utmd Anderson Cancer Center | USA | 72 | 2.869 |
| 9 | Capital Medical University | China | 70 | 2.789 |
| 10 | Pennsylvania Commonwealth System of Higher Education | USA | 70 | 2.789 |

**Table 5.**
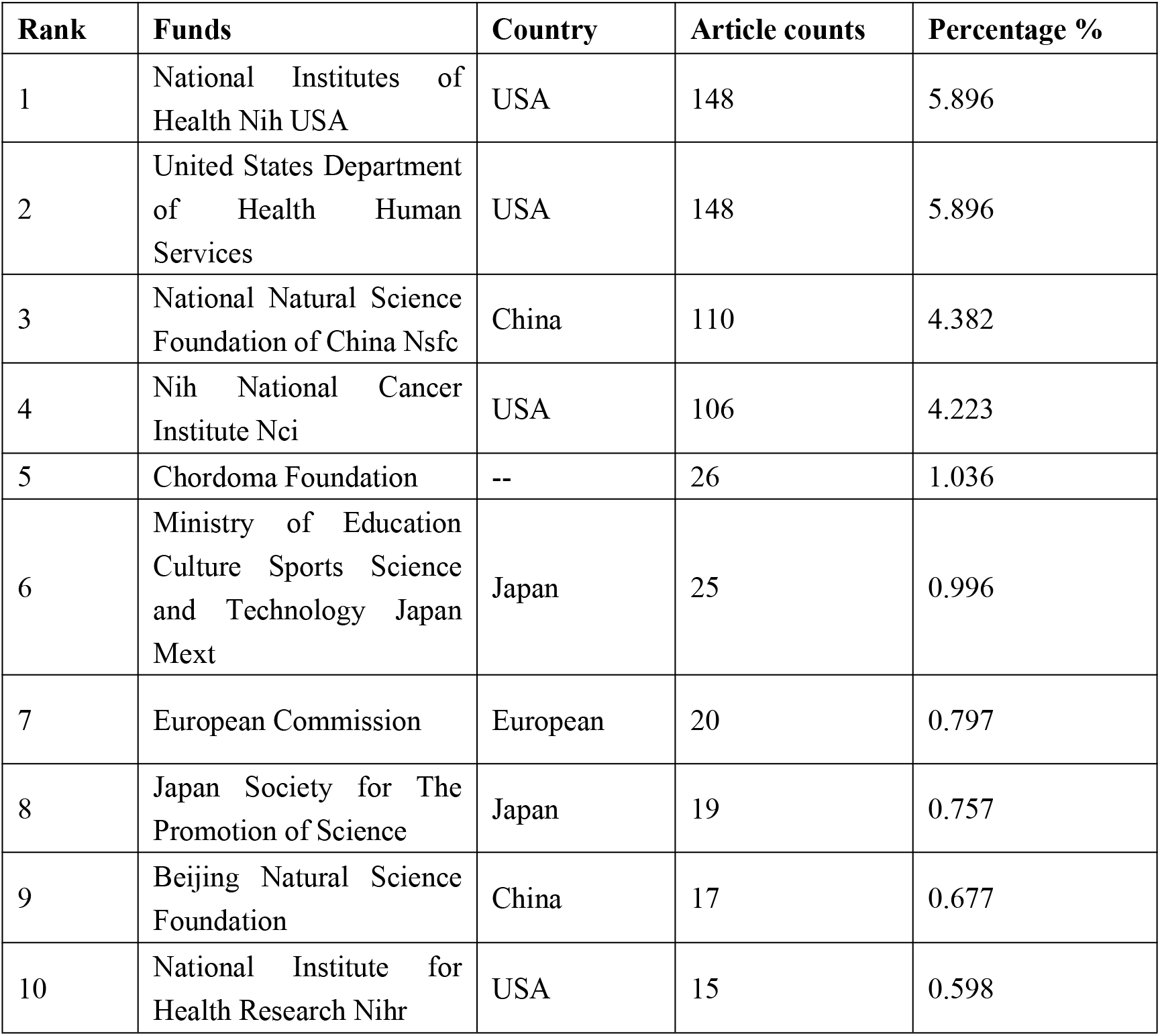
The top 10 funds with most publications related to chordoma research from 1991-2021.

### Bibliographic Coupling Analysis

Bibliographic co-citation analysis was also employed to investigate the interrelatedness among documents. Figure 4A depicts the results obtained using VOS viewer for papers originating from 37 countries, with a minimum of 5 documents per country selected for analysis. The USA (total link strength = 793460 times), China (total link strength = 336723 times), Italy (total link strength = 273038 times), Japan (total link strength = 202155 times), and England (total link strength = 199206 times) emerged as the top 5 countries with the highest total link strength in this field.

**Figure 4.**
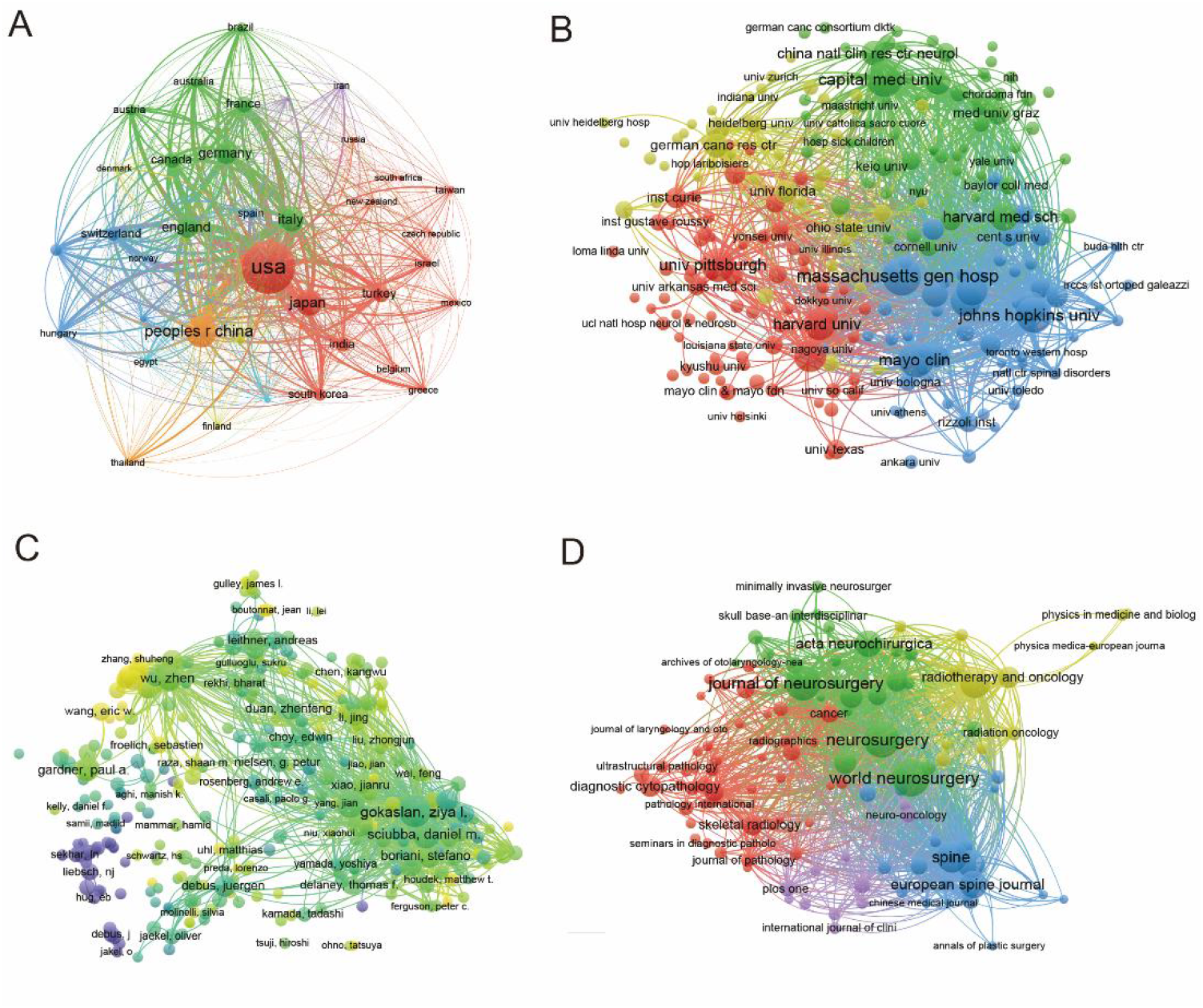
(A) Mapping of the 37 countries related to chordoma research from 1991-2021. (B) Mapping of the 233 institutions related to chordoma research from 1991-2021. (C) Density visualization of the 320 authors related to chordoma research from 1991-2021. (D) Mapping of the 114 identified journals related to chordoma a research from 1991-2021. The line between different points represents that the countries/institutions/journals/authors had establish a similarity relationship. The thicker the line, the closer the link between the journals/institutions/countries/authors. Yellow means appearing more frequently, however green means appearing less frequently.

For institutions, publications meeting the criteria of a maximum of 25 organizations per document and a minimum of 5 documents per organization were selected for analysis using VOS viewer, as shown in Figure 4B. The top 5 institutions with the greatest total link strength were Massachusetts General Hospital (total link strength = 136035 times), Johns Hopkins University (total link strength = 133354 times), Mayo Clinic (total link strength = 112860 times), Memorial Sloan Kettering Cancer Center (total link strength = 99934 times), and Capital Medical University (total link strength = 97455 times).

Regarding authors, publications with a minimum of 5 documents per author were selected, resulting in the analysis of 320 authors using VOS viewer. Figure 4C highlights the top 5 productive authors, namely Ziya L. Gokaslan (total link strength = 141590 times), Daniel M. Sciubba (total link strength = 95895 times), Francis Hornicek (total link strength = 93726 times), Joseph H. Schwab (total link strength = 90269 times), and Stefano Boriani (total link strength = 67787 times).

Furthermore, we employed VOS viewer to investigate the journals that appeared in the total publications, resulting in the identification of 114 journals with a total link strength, as depicted in Figure 4D. The top 5 journals with the highest total link strength were Neurosurgery (impact factor = 5.315, 2021, total link strength = 67771 times), World Neurosurgery (IF = 2.210, 2021, total link strength = 67157 times), Journal of Neurosurgery (IF = 5.408, 2021, total link strength = 67144 times), Spine (IF = 3.241, 2021, total link strength = 49733 times), and International Journal of Radiation Oncology (IF = 5.859, 2021, total link strength = 46021 times).

### Co-Citation Analysis of leading authors

The analysis of bibliographic coupling relied on the interrelationship between items based on their cocitation frequency. The ten most frequently cocited authors, as presented in **Table 6**, were found to have a total citation count exceeding 200. Among them, four were from the USA, two from Italy, and one each from Sweden, Switzerland, France, and Japan. Notably, Stacchiotti, S from Italy had the highest total citation count of 611, followed by Boriani, S (total citations: 600), and McMaster, ML (total citations: 456).

**Table 6.** The top 10 co-cited authors on chordoma research from 1991-2021.

| **Rank** | **Co-cited Authors** | **Country** | **Total citations** |
| --- | --- | --- | --- |
| 1 | Stacchiotti, S | Italy | 611 |
| 2 | Boriani, S | Italy | 600 |
| 3 | McMaster, Ml | USA | 456 |
| 4 | Bergh, P | Sweden | 352 |
| 5 | Sundaresan, N | USA | 326 |
| 6 | Hug, Eb | Switzerland | 304 |
| 7 | Walcott, Bp | USA | 277 |
| 8 | Noel, G | France | 255 |
| 9 | Heffelfinger, Mj | USA | 249 |
| 10 | Yamaguchi, T | Japan | 247 |

Using VOSviewer, a total of 616 authors with at least 20 publications were analyzed, and the top five authors with the highest total link strength were Sacchetti (total link strength = 15971 times), Biriani (total link strength = 13219 times), McMaster (total link strength = 10458 times), Hug (total link strength = 8805 times), and Bergh, P (total link strength = 8289 times), as depicted in Figure 5A.

**Figure 5.**
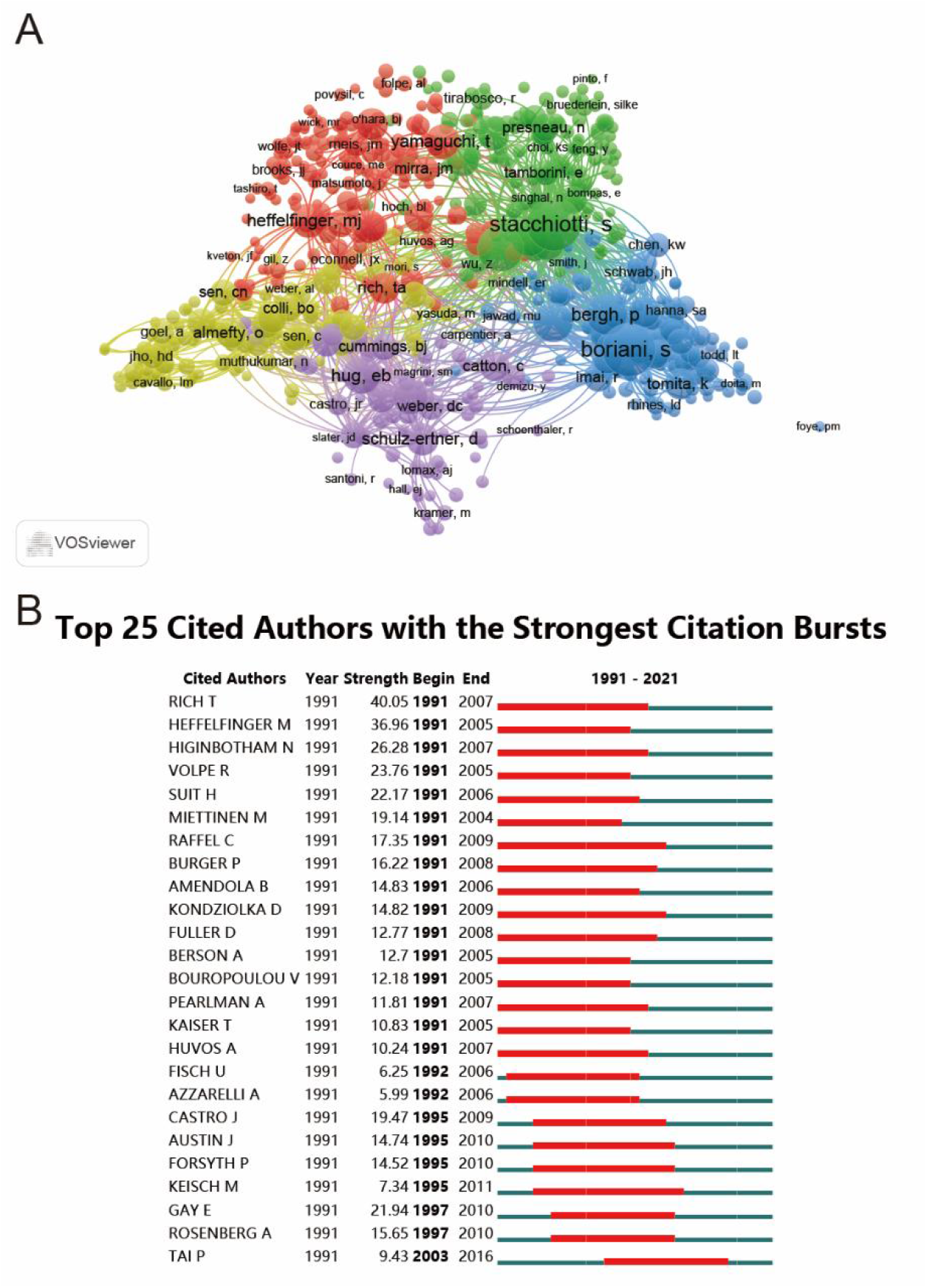
(A) Mapping of the co-cited authors related to chordoma research from 1991-2021. (The 616 points with different colors represent the 616 cited references.) (B) Top 25 authors with strongest citation bursts of publications related to chordoma research from 1991-2021. The point sizes represent the citation frequency. The line between different points indicates that they were cited in one paper. The shorter the line, the closer the link between two papers. The same color of the points represents the same research area they belong to.

Furthermore, we employed CiteSpace software to identify the burst of co-cited authors from 1991 to 2021, where the red line indicates the duration of the cited author’s burst, and the blue line represents the timeline (Figure 5B). The top 25 most intensely cited authors had a strength of more than 5. Rich T. had the most prominent burst strength (40.05), followed by Heffelfinger M. and Higinbotham N. (both with a strength of 36.96) starting from 1991 and ending in 2005 to 2007. Raffel C. and Kondziolka D. had the longest burst time of 18 years (1991 to 2009), while Tai P. had the most recent burst from 2003 to 2016, with a total linking strength of 9.43, possibly playing a leading role in this field.

### Co-Citation Analysis of leading journals

Utilizing VOSviewer, a cocitation analysis was conducted on the names of journals with a minimum of 20 citations. In Figure 6A, the total link strength displayed 432 journals, and the top 5 journals with the greatest total link strength were as follows: Journal of Neurosurgery (total link strength = 267764 times), Neurosurgery (total link strength = 217261 times), Laryngoscope (total link strength = 141793 times), International Journal of Radiation Oncology Biology Physics (total link strength = 132621 times), and World Neurosurgery (total link strength = 107519 times). Additionally, the top 10 cocited journals in this research area were determined, with Journal of Neurosurgery leading the way at No. 1 with a total of 4469 citations, followed by International Journal of Radiation Oncology Biology Physics and Neurosurgery with 3625 and 3579 citations, respectively (**Table 7**).

**Figure 6.**
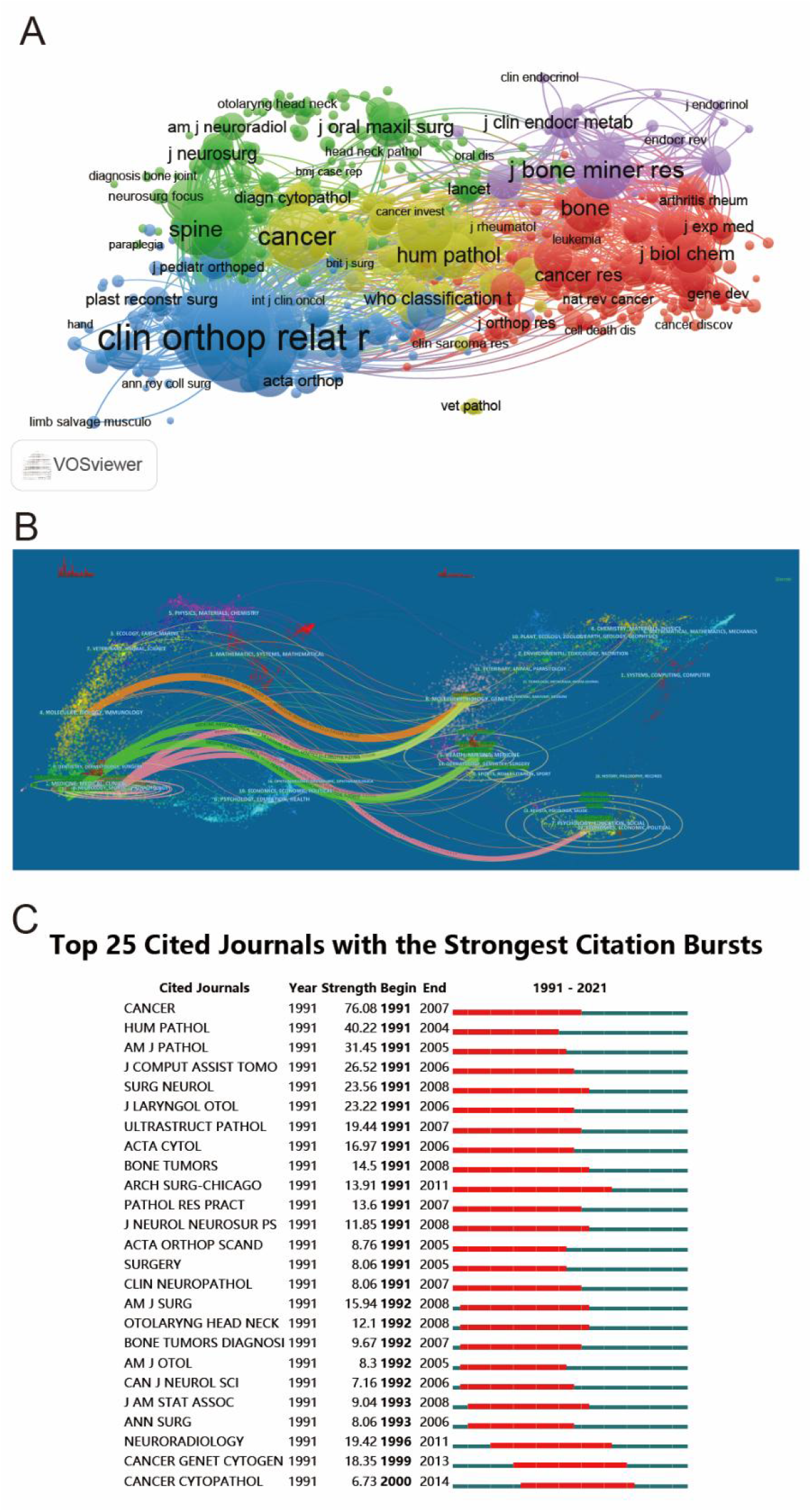
(A) Mapping of the co-cited journals related to chordoma research from 1991-2021. (The 432 points with different colors represent the 432 identified journals.) (B) The dual-map overlay of journals related to chordoma research from 1991-2021. (C) Top 25 journals with strongest citation bursts of publications related to chordoma research from 1991-2021. The point sizes represent the citation frequency. The line between different points indicates that they were cited in one paper. The shorter the line, the closer the link between two papers. The same color of the points represents the same research area they belong to.

**Table 7.** The top 10 co-cited journals related to chordoma research from 1991-2021.

| **Rank** | **Cited Journal** | **Citations** | **IF (2021)** |
| --- | --- | --- | --- |
| 1 | Journal of Neurosurgery | 4469 | 5.408 |
| 2 | International Journal of Radiation Oncology<br>Biology Physics | 3625 | 6.686 |
| 3 | Neurosurgery | 3579 | 5.315 |
| 4 | Spine | 2681 | 3.241 |
| 5 | Cancer | 1945 | 6.921 |
| 6 | American Journal of Surgical Pathology | 1473 | 6.298 |
| 7 | Journal of Bone Joint Surgery | 1090 | -- |
| 8 | Clinical Orthopedics and Related Research | 1053 | 4.755 |
| 9 | Journal of Neurosurgery-Spine | 957 | 3.67 |
| 10 | Journal of Clinical Oncology | 861 | 50.717 |

Moreover, the dual-map overlay of the journals was utilized to analyze the distribution of citing and cited journals, as illustrated in Figure 6B. The left points represent citing journals, the right ones represent cited journals, and the colored paths depict the citation relationship. In general, six colored primary citation pathways were found in Figure 6B. The orange pathway indicates that articles published in journals in the fields of molecular/biology/immunology were mostly cited by articles published in the fields of molecular/biology/genetics. The green lines were divided into two roads, with one road denoting that research published in the fields of medicine/medical/clinical was mainly cited by research in the fields of molecular/biology/genetics, while the other indicated that research in the fields of medicine/medical/clinical was primarily cited by research published in the fields of health/nursing/medicine. Furthermore, the pink line was divided into three lines, suggesting that articles published in the field of neurology/sports/ophthalmology were mainly cited by molecular/biology/genetics, health/nursing/medicine, and psychology/education/social.

Lastly, the top 25 cited journals with the strongest citation bursts were analyzed using CiteSpace, as displayed in Figure 6C. The most intense journals were Cancers (strength = 76.08), followed by Human Pathology (strength = 40.22) and American Journal of Pathology (strength = 40.22). Additionally, Archives of Surgery had the longest burst time of 20 years from 1991 to 2011. Notably, Cancer Cytopathology was the most recent burst journal from 2000 to 2014.

### Co-Citation Analysis of leading references

The utilization of reference cocitation analysis represents a valuable technique for elucidating the interconnectedness of items within a particular research field by assessing the overall number of citations and tracking the forefront of its evolution[27]. Additionally, VOSviewer was employed to analyze 533 references, which were chosen based on a minimum of 20 documents per cited reference, as depicted in Figure 7A. The larger nodes signify that the journals and authors have been cited more frequently. As illustrated in Figure 7A, the top 5 publications with the most significant total link strength were McMaster ml,2001, cancer cause control, v12, p1 (total link strength = 8103 times); bergh p,2000, cancer-am cancer soc, v88, p2122 (total link strength = 5755 times); Walcott bp,2012, lancet Oncol, v13, pe69 (total link strength = 4676 times); Catton c,1996, radiotherapy Oncol, v41, p67 (total link strength = 4331 times); york Je,1999, neurosurgery, v44, p74 (total link strength = 4020 times).

**Figure 7.**
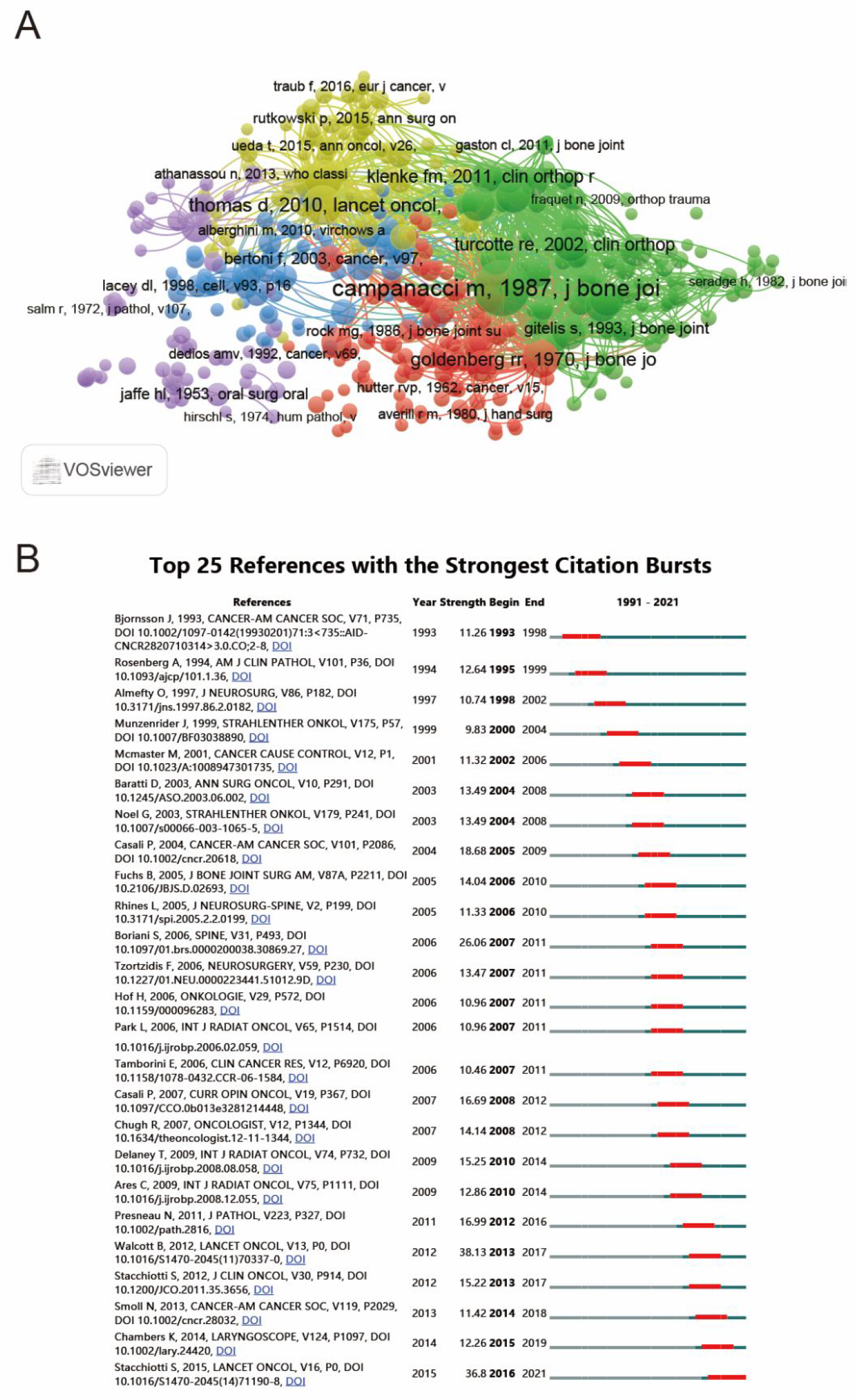
(A) Mapping of the co-cited references related to chordoma research from 1991-2021. (The 616 points with different colors represent the 616 cited references.) (B) Top 25 references with strongest citation bursts of publications related to chordoma research from 1991-2021. The point sizes represent the citation frequency. The line between different points indicates that they were cited in one paper. The shorter the line, the closer the link between two papers. The same color of the points represents the same research area they belong to.

**Table 8** presents the top 5 documents with the most citations, including Chordoma: incidence and survival patterns in the United States, 1973-1995 (total citation: 607), Endoscopic endonasal transsphenoidal surgery: Experience with 50 patients (total citation: 535), Bone Cancers (total citation: 418), Prognostic factors in chordoma of the sacrum and mobile spine - A study of 39 patients (total citation: 397), and Chordoma: current concepts, management, and future directions (total citation: 395). Of note, the document entitled Chordoma: current concepts, management, and future directions demonstrated the largest average citations, suggesting that it may have garnered more attention from researchers.

**Table 8.**
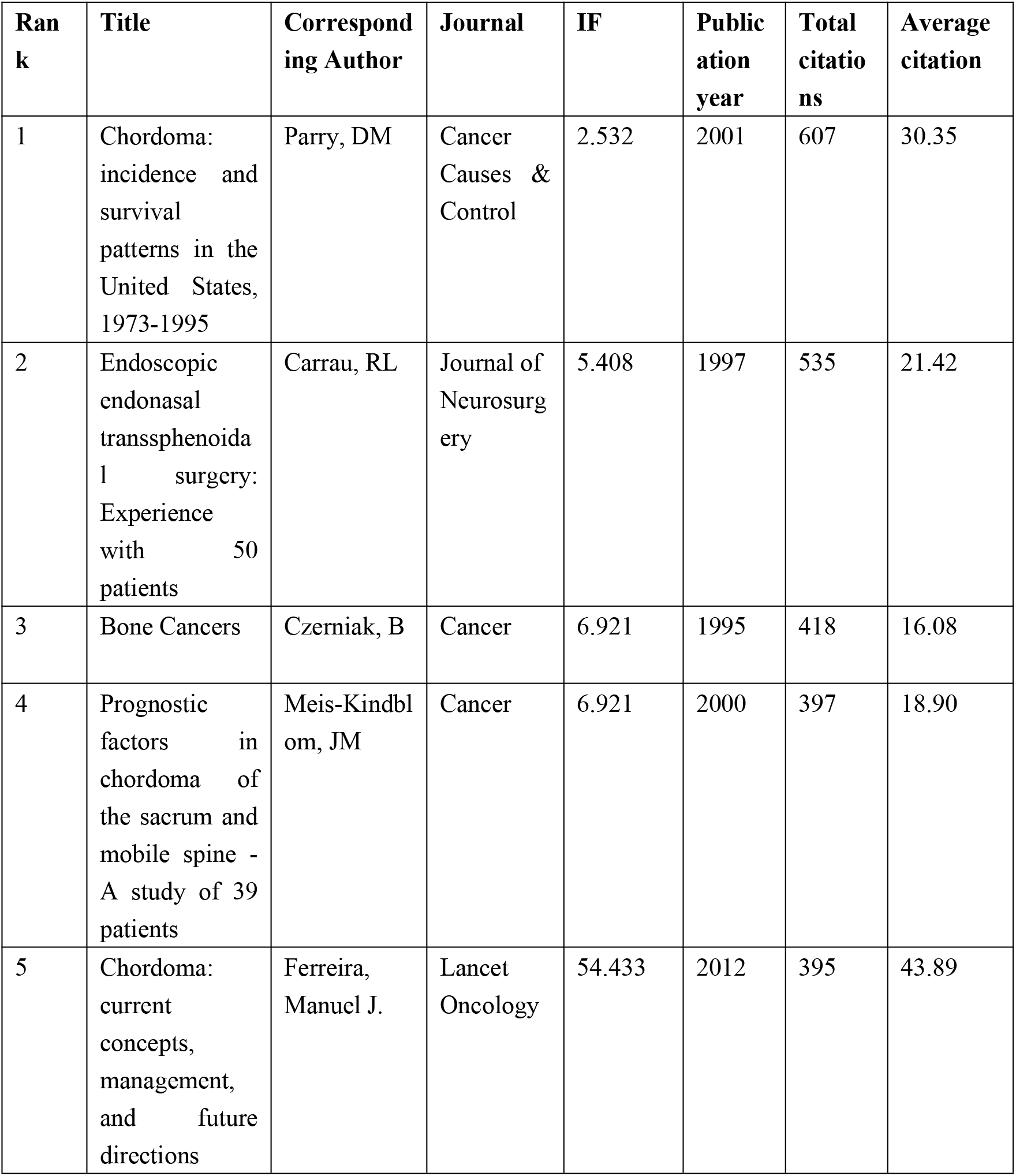
The top 5 documents with the most citations in chordoma research from 1991-2021.

| **Ran**<br>**k** | **Title** | **Correspond**<br>**ing Author** | **Journal** | **IF** | **Public**<br>**ation**<br>**year** | **Total**<br>**citatio**<br>**ns** | **Average**<br>**citation** |
| --- | --- | --- | --- | --- | --- | --- | --- |
| 1 | Chordoma: incidence and survival patterns in the United States, 1973-1995 | Parry, DM | Cancer Causes & Control | 2.532 | 2001 | 607 | 30.35 |
| 2 | Endoscopic endonasal transsphenoidal surgery: Experience with 50 patients | Carrau, RL | Journal of Neurosurgery | 5.408 | 1997 | 535 | 21.42 |
| 3 | Bone Cancers | Czerniak, B | Cancer | 6.921 | 1995 | 418 | 16.08 |
| 4 | Prognostic factors in chordoma of the sacrum and mobile spine - A study of 39 patients | Meis-Kindblom, JM | Cancer | 6.921 | 2000 | 397 | 18.90 |
| 5 | Chordoma: current concepts, management, and future directions | Ferreira, Manuel J. | Lancet Oncology | 54.433 | 2012 | 395 | 43.89 |

Furthermore, the 25 references with the strongest citation bursts were investigated by CiteSpace and are displayed in Figure 7B. The reference by Walcott B, 2012, LANCET ONCOL, V13, P0, DOI 10.1016/S1470-2045(11)70337-0 possessed the highest citation strength of 38.13, followed by Stacchiotti S, 2015, LANCET ONCOL, V16, P0, DOI 10.1016/S1470-2045(14)71190-8 (strength: 36.8) and Boriani S, 2006, SPINE, V31, P493, DOI 10.1097/01.brs.0000200038.30869.27 (strength: 26.06). The reference with the most recent burst time was Stacchiotti S, 2015, LANCET ONCOL, V16, P0, DOI 10.1016/S1470-2045(14)71190-8 from 2016 to 2021, indicating the latest area of research interest.

### Co-authorship Analysis of author, country, and institutions

In Figure 8A, an overlay visualization map was employed to assess the relatedness of items based on the total number of coauthored papers using VOSviewer. As depicted in Figure 8A, 320 authors with more than 5 documents were selected and analyzed. The top 5 authors with the highest total link strength were gokaslan, Ziya l. (total link strength = 221 times), scuba, daniel m. (total link strength = 181 times), Hornick, franchise. (total link strength = 169 times), Wu, Zhen (total link strength = 159 times), and wang, Liang (total link strength = 150 times).

**Figure 8.**
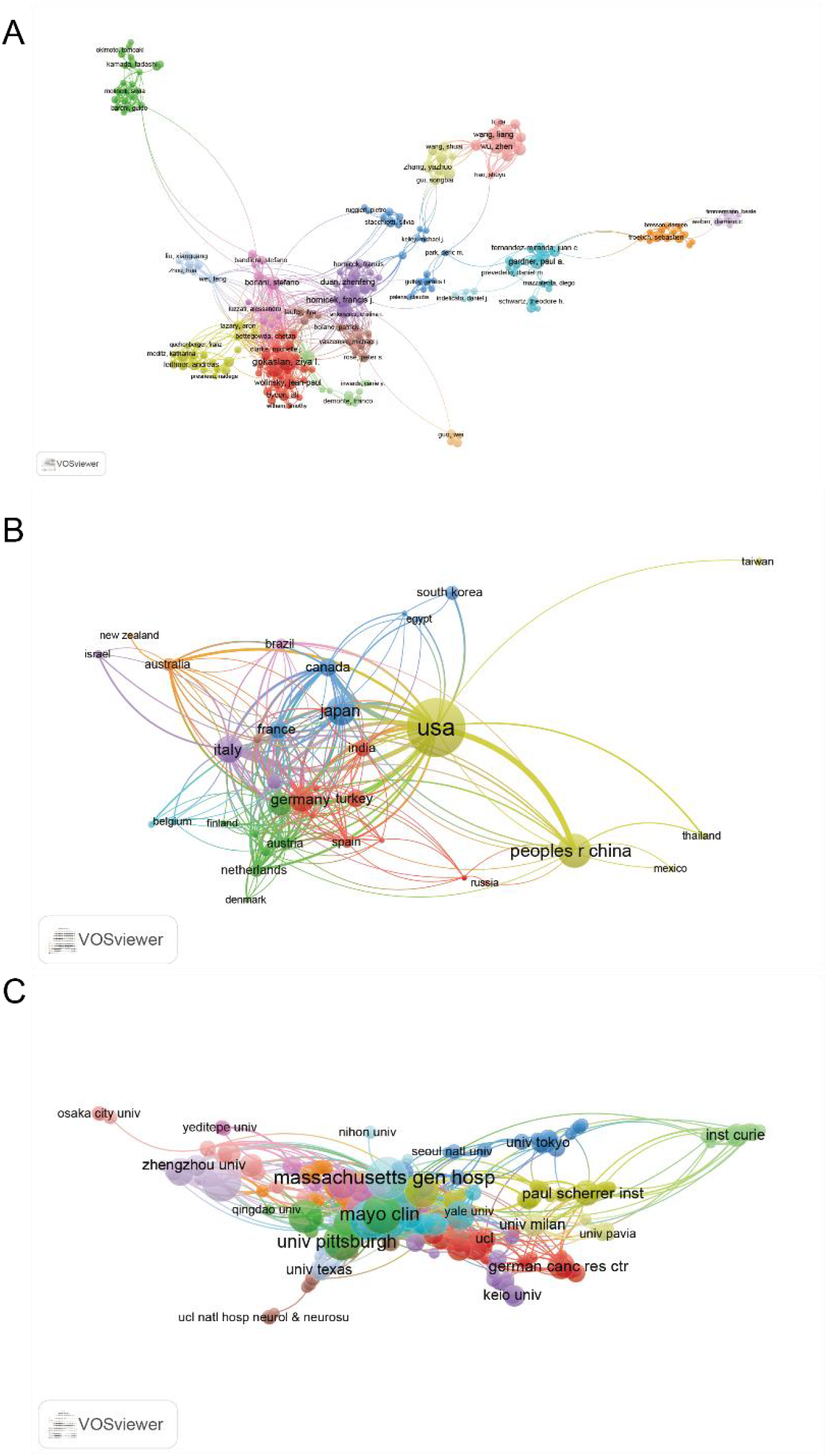
(A) Mapping of the 209-author co-authorship analysis related to chordoma research from 1991-2021. (B) Mapping of the 37-country co-authorship analysis related to chordoma research from 1991-2021. (C) Mapping of the 223-organizations co-authorship analysis related to chordoma research from 1991-2021. The line between two points represents the establishment of collaboration within two authors/institutions/countries, where the thickness indicates the collaboration strength between the two authors/institutions/countries. Yellow means appearing more frequently, however green means appearing less frequently.

To further investigate the coauthorship patterns, VOSviewer was also used to analyze 37 countries with over 5 papers, as illustrated in Figure 8B. The USA had the most significant total link strength (total link strength = 358 times), followed by Italy (total link strength = 198 times), England (total link strength = 155 times), Germany (total link strength = 148 times), and Canada (total link strength = 129 times).

Furthermore, 233 institutions with more than 5 documents were analyzed using VOSviewer, as shown in Figure 8C. The top 5 institutions with the highest total link strength were Johns Hopkins Univ (total link strength = 169 times), Univ Toronto (total link strength = 139 times), Univ Texas Md Anderson Canc Ctr (total link strength = 130 times), Massachusetts Gen Hosp (total link strength = 126 times), and Univ British Columbia (total link strength = 118 times).

### Co-occurrence Analysis of Keywords

To begin with, we employed CiteSpace to conduct a co-occurrence cluster analysis of keywords, with the aim of investigating the current research frontiers. Our analysis revealed 12 distinct clusters, including expression (cluster 0), proton therapy (cluster 1), en bloc resection (cluster 2), immunohistochemistry (cluster 3), pituitary adenoma (cluster 4), benign notochordal (cluster 5), endoscopic endonasal surgery (cluster 6), gliosarcoman (cluster 7), myxoid adrenocortical carcinoma (cluster 8), congenital cyst (cluster 11), chloroquine (cluster 12), and epidural tumor (cluster 13) (Figure 9A). Additionally, we plotted the dynamic evolution of the keyword clusters over time using CiteSpace, which is presented in Figure 9B. Our analysis indicated that clusters 0 to 5, 7, and 12 were former hotspots, while clusters 6 and 11 were mid-period hotspots and cluster 8 was a present research hotspot.

**Figure 9.**
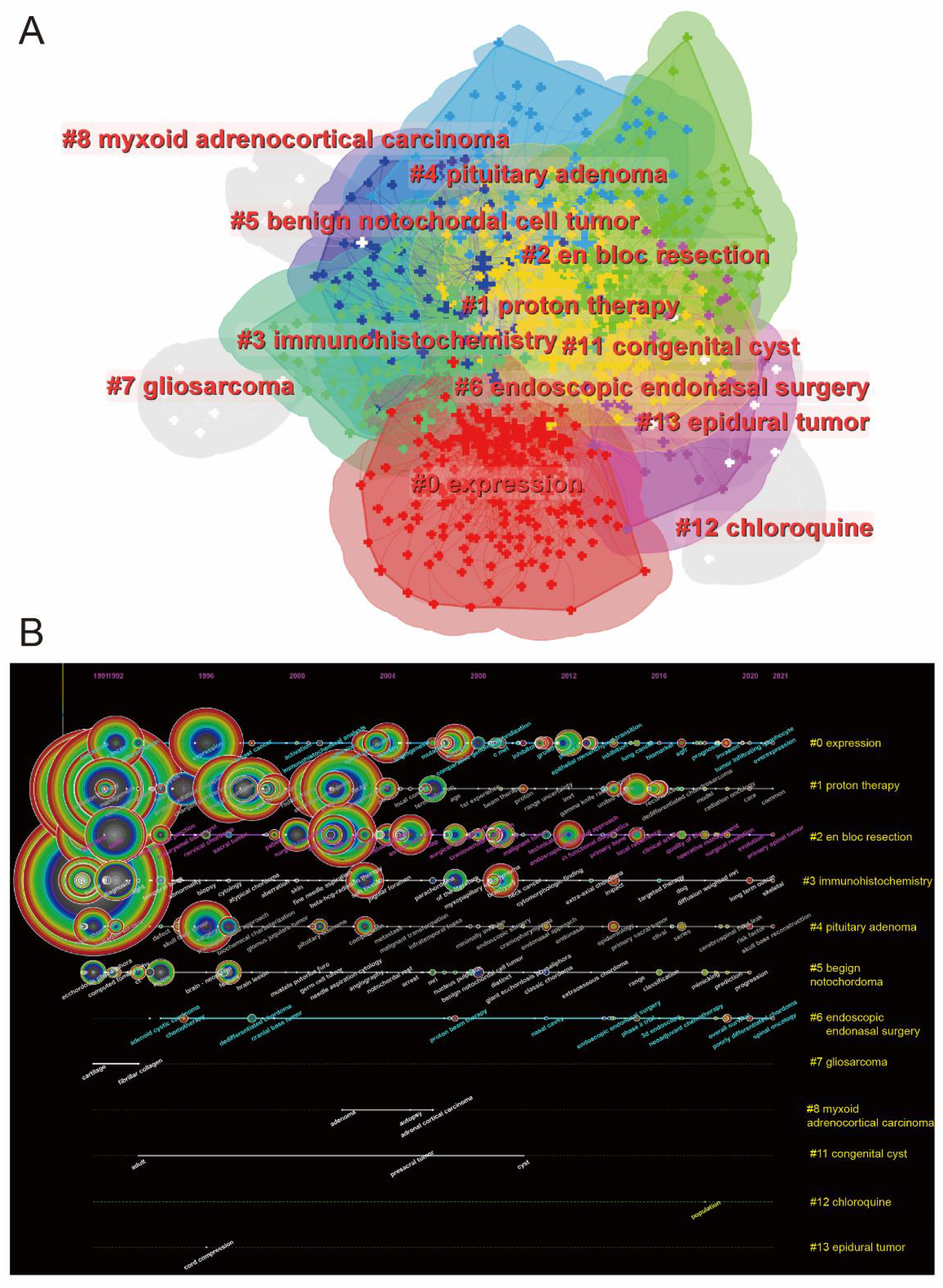
(A) Clustering map of occurrence analysis of keywords based on CiteSpace. (B) Keyword timeline visualization from CiteSpace.

In bibliometrics, co-occurrence analysis of keywords is a widely-used method to identify key research areas and topics, and it plays a crucial role in monitoring advancements in scientific research. To identify hot research areas in chordoma research, we performed a co-occurrence analysis of the 744 identified keywords using VOS viewer. The results were generally divided into five clusters, including cluster 1: radiotherapy study (purple), cluster 2: basic and clinical trial research (green), cluster 3: diagnostic study (red), cluster 4: surgical approach research (blue), and cluster 5: prognostic study (yellow), as depicted in **Figure 10A**. Our analysis revealed that the primary research topics in chordoma research thus far include chondrosarcoma, radiation therapy, and radiotherapy in the “radiotherapy study” cluster, expression, cancer, and survival in the “basic and clinical trial research” cluster, tumors, tumor, and immunohistochemistry in the “diagnostic study” cluster, management, skull base, and surgery in the “surgical approach research” cluster, and prognosis factors, spine, and experience in the “prognostic study” cluster. Additionally, **Figure 10B** shows the trends of chordoma research colored by VOS viewer, indicating that the research hotspots have shifted from clusters 3 and 4 to clusters 1, 2, and 5, which may indicate changes in future research directions.

**Figure 10.**
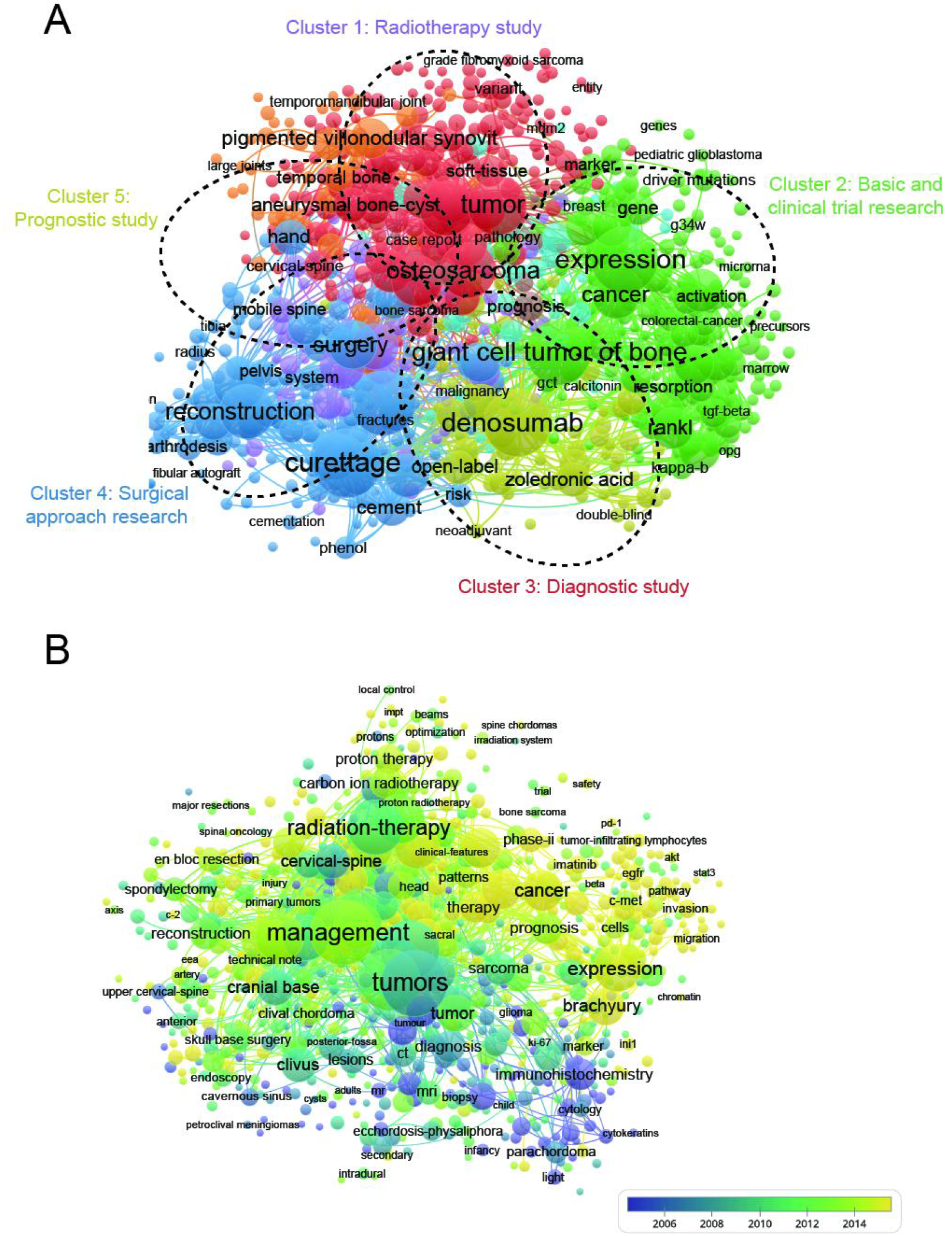
(A) Mapping of keywords in the research related to chordoma research from 1991-2021; the frequency is represented by point size and the keywords of research fields are divided into six clusters: rheumatoid arthritis research (red), clinical symptoms (green), regeneration research (yellow), mechanism (dark blue), pathological features (dark brown), and surgery research (baby blue). (B) Distribution of keywords according to the mean frequency of appearance; keywords in yellow appeared later than those in blue.

To further explore the research hotspots, we conducted a burst analysis of keywords using CiteSpace, with a minimum burst duration of 2 years. Figure 11 displays the top 25 keywords with the strongest citation bursts related to chordoma research. Our analysis indicated that the keyword “skull” had the largest burst strength of 19.13, followed by “base” (strength: 18.15) and “chondroid chordoma” (strength: 15.62). The keyword with the longest burst time was “mr” from 1991 to 2010, and the latest burst keyword was “sacrectomy” from 2009 to 2018.

**Figure 11.**
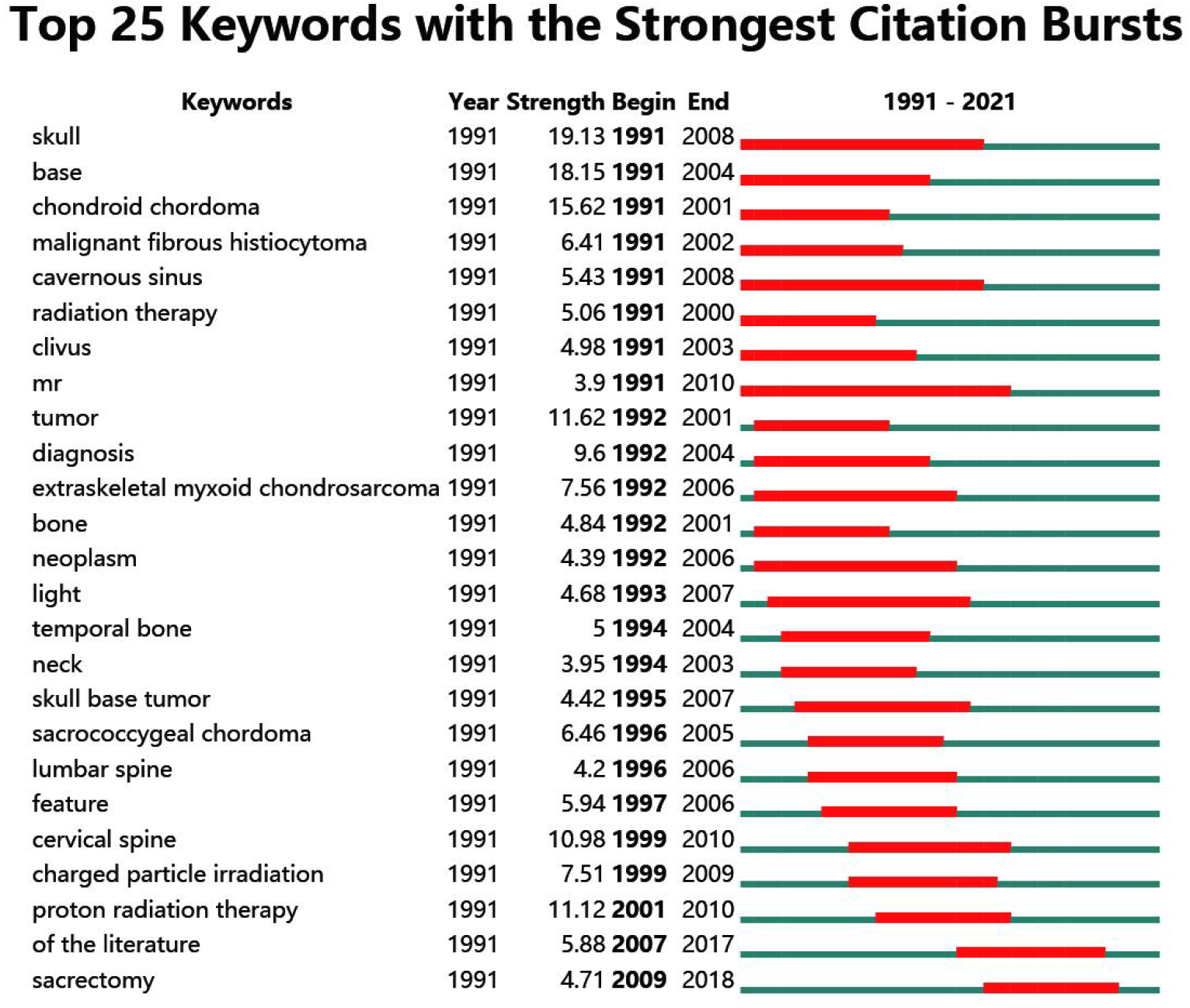
Top 25 keywords with the strongest citation bursts of publications related to chordoma research from 1991-2021.

## Discussion

In the current investigation, we conducted a bibliometric and visualization analysis to examine the recent research landscape of chordoma from 1991 to 2021 and prognosticate future developments. We explored the spatiotemporal distribution of publications, as well as the contributions of journals, authors, institutions, and countries, research areas, keywords, and potential research directions and hotspots to elucidate the evolution of readers’ thought processes and research transformations.

### The Trend Overview of Global Publications of chordoma

Since Virchow’s initial observation of a small mucinous excrescence on the clivus in 1846, numerous researchers have independently described similar structures[28–31]. In 1894, Ribbert corroborated Muller’s theory through animal experimentation and coined the term “chordoma”[32]. Over the past few decades, significant strides have been made in chordoma diagnosis and treatment through substantial research efforts[1, 2, 33]. Currently, gross total resection, when safely feasible, is the primary treatment and consistent prognostic factor. However, the eradication of recurrent chordoma is nearly impossible, leading surgeons to aim for ablative resection with wide margins. Thus, a significant hurdle in chordoma research is the development of basic research and effective treatments.

Our findings indicate a marked rise in the number of publications per year from January 1, 1991, to December 31, 2021. Additionally, the RRI has marginally increased in recent years. In our analysis, approximately 63 countries and 2056 institutions have published papers in the field of chordoma. Notably, the United States has contributed the highest number of papers (1000, 39.84%) compared to China (333, 13.27%), Japan (256, 10.20%), Italy (211, 8.41%), and Germany (178, 7.09%).

Among the academic institutions, Harvard University (151 publications), League of European Research Universities (Leru) (150 publications), and Massachusetts General Hospital (121 publications) have actively contributed to the research forefront. Consequently, this evidence collectively suggests that collaborative, in-depth studies could play a pivotal role in chordoma research, enabling researchers to publish high-quality papers in the future.

### Quality and Status of Global Publications

Recently, bibliometric studies have increasingly utilized total citations, per citations, and H-index as critical parameters to measure the academic impact and quality of different countries. As displayed in Figures 2 and 3, the USA leads in this field with the most extensive total citations and largest H-index, indicating the country’s highly productive and preeminent position. With the most elite researchers and institutions globally, the USA’s leadership in chordoma research is indisputable. Notably, Sweden, Switzerland, and England rank highest in average citations, surpassing the USA.

Italy, though ranking fourth in the number of publications, has made considerable strides in total citations, ranking second and third in H-index. China, on the other hand, ranks second in total publications but fares poorly in total citations, average citations, and H-index, potentially lagging behind the USA in the coming years. Such inconsistencies between quantity and quality of publications in China necessitate significant efforts by the Chinese academic evaluation systems (CAESs) in this field.

In terms of journals, World Neurosurgery, Neurosurgery, and Journal of Neurosurgery published the most papers, with high impact factors in chordoma research. Notably, the top 5 journals published over 50 papers, making them prime candidates for future high-quality research. Regarding institutions, the top 5 institutes have significantly contributed to chordoma research, aligning with the global publications produced by the top 5 countries (Table 4). Moreover, the top 10 institutes are from the top 5 countries, highlighting the critical role of first-class institutions in improving a country’s academic research ranking. The top-ranked authors, predominantly Americans and affiliated with Harvard University, along with generous funding from the USA National Institutes of Health (NIH), have played an integral role in advancing chordoma research. These top-ranked authors, listed in Table 3, were early entrants in chordoma research, indicating their keen attention to new advancements. Previous reports have employed bibliometric research methods to identify research hotspots.

In the field of bibliometrics, coupling analysis was carried out to establish a comparable relationship among the chosen articles from the country, institution, journal, and author selected for our study. Figure 4 demonstrates that the USA was the most closely related country, Massachusetts Gen Hosp was the most strongly associated institution, Gokaslan and Ziya L. were the most related authors, and Neurosurgery was the most connected journal. Additionally, we conducted a co-citation analysis based on authors, journals, and references to investigate the impacts of publications by analyzing the total number of citations (Figure 5-7, Table 6-8). The leading author with the highest number of citations and total link strength (15971) was Stacchiotti, S, while Rich T. was the most notable author with the strongest citation burst (Figure 5, Table 6). We also explored the co-cited journals associated with chordoma research from 1991-2021, and the results indicated that the Journal of Neurosurgery had the most citations and the largest total link strength, while Cancers had the journals with the strongest citation burst (Figure 6, **Table 7**). Furthermore, our results in Figure 7 and Table 8 revealed that the most impactful studies about chordoma research had the highest citation frequency in this field. Interestingly, a co-creation analysis of journals can determine which journals have made the most significant contributions in this field. As shown in Table 1 and Table 3, Cacciotti and J Neurosurg may be the top authors or journals with the highest citation frequency, respectively. Co-authorship analysis can reflect the connections between authors, countries, and institutions. Therefore, authors/countries/institutions with a high total link strength should work collaboratively. Based on the evidence from these findings, we could suggest that cross-cooperation among different authors/countries/institutions in future research can significantly enhance communication and productivity and ultimately improve the research level in the specific subject.

### Research Focus on chordoma Research

The analysis of keyword co-occurrence has unveiled the developing trends and pivotal areas in chordoma research. In this study, we constructed a map of occurrence networks based on the identification of keywords in the titles/abstracts of all included publications. Initially, we categorized the keywords into 12 clusters and further analyzed the citation bursts and time characteristics of each cluster. As time progressed, the research focus has evolved. As presented in Figure 9, clusters 0 to 5, 7, and 12 were the former hotspots, clusters 6 and 11 were the mid-period hotspots, and cluster 8 is currently the hotspot. Additionally, we explored the strongest citation bursts and found that the skull has the largest burst strength, which could be attributed to the prevalence of chordoma in this area (Figure 11). Figure 10A depicts the 5 primary research trends, which can be classified into 5 clusters: radiotherapy study (purple), basic and clinical trial research (green), diagnostic study (red), surgical approach research (blue), and prognostic study (yellow). These results not only conform to the critical hotspots in the field of chordoma research but also forecast the future directions of research as follows:

I. Radiotherapy study: The co-occurrence analysis of keywords has identified “chondrosarcoma,” “radiation-therapy,” and “radiotherapy” as important research hotspots that require further investigation. Chondrosarcoma, which mainly occurs at the skull base, exhibits radiological features and clinical presentations similar to those of chordoma. However, chondrosarcoma has a better prognosis, with a median projected survival over 20 years[34]. Although radiotherapy has been considered an adjuvant treatment and has not been statistically shown to improve survival in many studies, the reason for the failure of radiotherapy treatment may be due to the underrepresentation of contemporary radiation treatment modalities[35]. Importantly, the administration of radiotherapy could be customized according to the type, method, and total dose of radiation, which warrants further investigation. For instance, in a study with surgical resection of chordomas, patients were stratified into three groups: surgical resection alone, therapeutic radiotherapy alone, and surgical resection plus therapeutic radiotherapy. The authors concluded that chordoma patients with positive margins exhibited improved survival after adjuvant radiotherapy, with a cumulative dose >65 Gy[35]. Recently, carbon ion radiotherapy (CIRT) has emerged as an alternative treatment when surgery is not preferred. Yolcu et al. compared the oncologic outcomes and treatment-related toxicity of CIRT with those of patients undergoing en bloc resection. They found that CIRT can be applied to patients without surgery or older patients with high-performance status and may also provide benefits for patients with margin-positive resection[36]. These studies shed light on the potential benefits of adjuvant radiotherapy for chordoma patients and emphasize the need for further exploration.
II. Fundamental and Clinical Trial Investigation: A primary focus of fundamental research involves exploring the in vitro gene expression of chordoma. At present, in vitro chordoma models are still scarce, with only five cell lines (U-CH1, U-CH2, MUG-Chor, JHC7, and UM-Chor1) approved for investigation[37]. In terms of gene expression related to tumor progression, chordomas have been found to express estrogen receptor α and progesterone receptor β[38]. Another potential therapeutic target is the programmed death-ligand (PD-1), which is expressed in tumor-infiltrating lymphocytes (TILs) and may be linked to poor chordoma prognosis[39]. The prevalence of TILs and chordoma progression were also highly correlated with PD-1 expression[39, 40]. Another critical aspect of clinical research is patient survival. Notably, Vanderheijden, Cas, et al. conducted a systematic review of 78 original studies examining expressed factors related to chordoma survival. The results identified 26 factors (such as Y box binding protein 1, Src-associated in mitosis, and extracellular signal-regulated kinase) with negative impacts, while 6 factors (such as miR-1237-3p, phosphorylated BAD, and CD8/Foxp3) exhibited positive impacts on patient survival[41]. These expressed factors seem to play a vital role in chordoma pathophysiology and merit further in-depth investigation in the future.
III. Diagnostic investigation: Urgent demand for personalized diagnoses of chordoma is imperative for precision medicine. Chordoma can be positively expressed for specific keratins (cytokeratin 8, CK18, and CK19), epithelial membrane antigen (EMA), and S100 protein, while negatively expressed for CK7 and CK20[11, 12, 41]. Immunohistochemistry plays a vital role in chordoma diagnosis, and numerous efforts have been made to explore novel, specific immunohistochemical markers for chordoma in recent years. For example, brachyury, a T-box transcription factor encoded by the TBXT gene on chromosome 6, can be applied as a highly sensitive and specific marker for chordoma diagnosis[11]. In addition, investigating the underlying molecular pathology of chordoma is a critical research direction for differential diagnosis. For instance, Zhang, Y et al. conducted an in vitro analysis and successfully identified that miR-608 and miR-34a are expressed at low levels in most chordoma cells and inversely correlate with epidermal growth factor receptor gene (EGFR) and mesenchymal-epithelial transition factor (MET) levels[42].
IV. Surgical strategy research: Despite complete en bloc resection of chordoma with clear margins and postoperative radiotherapy having been proven to result in the longest survival in most studies, the prognosis for chordoma remains poor[43]. Hence, advanced management of the chordoma perioperative period, involving surgeries and postoperative radiotherapy, is urgently needed. Notably, skull base chordomas present a wide range of surgical approaches, such as transsphenoidal, transaxillary, transnasal, high anterior cervical retropharyngeal, and transoral approaches[44–46], unlike chordomas of the mobile spine and sacrum. Skull base chordoma also differs from spine chordoma, as it cannot be resected en bloc[47]. A study by Stüer et al. demonstrated that subtotal or partial resection of skull base chordoma attenuates the occurrence of neurological deterioration, emphasizing the importance of integrated management, such as less aggressive operative procedures and optional radiotherapy[48].
V. Prognostic investigation: Multiple prognostic factors have been analyzed for chordoma prognosis. Seven studies have focused on the vertebral column[49–55]. The most common adverse prognostic factors within these studies were as follows: larger tumor, older age, inadequate surgical margins, and nonen bloc resection (compared to en bloc resection). However, due to patient group differences and various methodologies, it is not practical to conclude prognostic factors in general[56]. In addition to the common prognostic factors mentioned above, other frequently mentioned prognostic factors include adjuvant radiation therapy or a lower dose of radiation therapy. Regarding the current status of chordoma prognostic factor studies, we suggest that future research should focus on designing more systematic prospective studies. Thus, clinicians can provide better medical experiences and treatment for their patients.

### Limitations

There exist some limitations that need to be acknowledged. Firstly, as the studies were collected solely from WoSCC, it is possible that publication bias may have resulted from the exclusion of databases such as PubMed, Cochrane, and Embase. Secondly, due to the omission of non-English publications, some relevant articles may have been left out. Thirdly, the accuracy of bibliometric tools can be affected by issues like journal name changes, which could have influenced the final results to some degree. Additionally, since new studies are constantly being published, some crucial and groundbreaking research may have been overlooked. Therefore, we hope that more comprehensive analysis will be conducted in future studies.

## Conclusion

In conclusion, chordoma is a rare and malignant tumor that originates from remnants of the notochord. Recent years have seen significant development in chordoma research worldwide. This study is the first bibliometric analysis to provide a comprehensive and systematic overview of chordoma research trends over the past 30 years. The research hotspots in this field include radiotherapy studies, basic and clinical trial research, diagnostic studies, surgical approach research, and prognostic studies, highlighting the focus on radiotherapy efficacy, the relationship between basic and clinical trials, surgical strategies, and the importance of systematic treatment. It is expected that further cooperation among authors, institutions, and countries will drive advancements in chordoma research in the future.

## Acknowledgements

N/A.

